# Transferrin receptor-binding blood-brain barrier shuttle enhances brain delivery and efficacy of a therapeutic anti-Aβ antibody

**DOI:** 10.1101/2025.02.14.638216

**Authors:** Marta Ramos Vega, Henrik H. Hansen, Camilla Stampe Jensen, Evdoxia Alexiou, Martin R Madsen, Franziska Wichern, Jacob Lercke Skytte, Casper Graversen Salinas, Florence Sotty, Allan Jensen, Sandra Vergo, Jacob Hecksher-Sørensen

## Abstract

Transferrin receptor-1 (TfR1) transcytosis-mediated delivery of therapeutic monoclonal antibodies across the blood-brain barrier (BBB) is a promising concept in drug development for CNS disorders. We sought to investigate brain delivery and efficacy of Aducanumab (Adu), an anti-Aβ antibody, when fused to a mouse TfR1-binding Fab fragment as BBB shuttle (TfR1-Adu). Automated 3D light sheet fluorescence imaging coupled with computational analysis was applied to evaluate drug IgG distribution and plaque counts throughout the intact brain of transgenic APP/PS1 mice. TfR1-Adu demonstrated enhanced brain delivery and more homogeneous distribution after both acute and chronic dosing in transgenic APP/PS1 mice compared with unmodified Adu. Also, importantly, only unmodified Adu showed perivascular labelling. While high-dose Adu promoted Aβ plaque depletion in multiple brain regions, similar plaque-clearing efficacy was achieved with a five-fold lower dose of TfR1-Adu. Furthermore, low-dose TfR1-Adu demonstrated greater capacity to reduce congophilic plaque burden. Collectively, these observations strongly support the applicability of TfR1-enabled BBB shuttle strategies to improve brain delivery and plaque-clearing efficacy while mitigating the risk of vascular-associated amyloid-related imaging abnormalities (ARIA) adverse effects associated with current Aβ immunotherapeutics.

## INTRODUCTION

Brain accumulation of amyloid β (Aβ) plaques is considered a key molecular driver of Alzheimer’s disease (AD). In recent years, humanized monoclonal antibodies (mAbs) targeting Aβ, including Aducanumab (Aduhelm), Lecanemab (Leqembi) and Donenamab (Kisunla), have received accelerated FDA-approval for early symptomatic AD, constituting landmark events in disease-modifying therapies for AD^1–3^. These three anti-Aβ mAbs represent the first wave of Aβ-targeted therapies, and several anti-Aβ immunotherapies are in clinical development for AD^4,5^. While these breakthrough immunotherapeutics have been demonstrated to reduce Aβ plaque load, benefits on slowing cognitive decline have so far been minimal or modest in AD patients^1,3,6^, and the durability of therapeutic effect is uncertain at this time until long-term longitudinal outcome studies have been completed. In addition, concerns have been raised regarding an increased risk of vascular-associated neurological complications, classified as amyloid-related imaging abnormalities (ARIA) in the forms of brain microhaemorrhages/hemosiderosis (ARIA-H) and edema/effusions (ARIA-E), which typically occur early in the course of anti-amyloid immune therapy^1,3,7^. Also, accelerated lateral ventricular enlargement is a recognized complication of current Aβ immunotherapies and presumed to be mechanistically linked to ARIA^8^. Both complications prompt serial MRI monitoring and may lead to treatment discontinuation^9^.

A significant limitation of antibody-based drug therapies is the highly restricted blood-brain barrier (BBB) penetration, a semi-permeable barrier encompassing multiple microvasculature-associated cell types such as endothelial cells, pericytes, astrocytes and microglia, which allows only passive diffusion of low-molecular weight molecules^10^. Accordingly, it is estimated that a minimal fraction (∼0.1%) of systemically administered antibodies reach the CNS ^11–13^. Achieving adequate brain delivery therefore poses a fundamental challenge in the development of more effective therapeutic antibodies for AD^14,15^. Shuttle antibodies binding to BBB endothelial transmembrane proteins have been intensely explored as molecular ‘Trojan horses’ which can carry a fused mAb across the BBB to access the brain parenchyma^15,16^. Among these, transferrin receptor-1 (TfR1) has gained significant attention due its high expression on brain endothelial cells, where it performs a critical role in cellular iron uptake through receptor-mediated endocytosis of iron-bound transferrin^17–20^. A major hurdle in assessing brain delivery of therapeutic mAbs is the complex 3D organization of the brain. To date, various TfR1 shuttles have been reported to enhance brain delivery and plaque clearing efficacy of humanized Aβ antibodies in mouse models of AD, evaluated using methods with limited spatial information and/or resolution including brain homogenate assays, conventional 2D histology, autoradiography, positron emission tomography (PET) or single-photon emission computed tomography (SPECT)^21–27^.

To further advance preclinical drug discovery for AD, brain distribution and therapeutic effects of Aβ antibody drug candidates should optimally be evaluated at single-plaque resolution on a brain-wide scale. Tissue clearing combined with light sheet fluorescence microscopy (LSFM) has emerged as a powerful tool in preclinical neuroscience and drug discovery, enabling unbiased 3D mapping of histological markers, drug effects and biodistribution in the intact mouse brain at cellular resolution^28–30^. While LSFM imaging has recently proven instrumental in preclinical AD research^31–35^, it remains to directly connect benefits on plaque burden with local brain exposure levels of Aβ-targeted biologics. Further 3D characterization could therefore help clarifying brain delivery and pharmacodynamics of anti-Aβ mAbs in mouse models of AD. To this end, we developed a high-throughput quantitative LSFM pipeline enabling mapping and quantification of anti-Aβ IgG distribution against plaque architecture across the entire mouse brain at single-region level. Using this platform, we profiled Aducanumab (Adu) fused with a mouse anti-TfR1 fragment (TfR1-Adu) in a standard transgenic APP/PS1 mouse model of AD. Our study indicates brain-wide delivery and efficacy of TfR1-Adu at significantly lower doses compared to unmodified Adu, opening new avenues for discovering TfR1-shuttled Aβ mAbs with increased brain parenchymal access, therapeutic efficacy and reduced risk for vascular-associated adverse effects in AD.

## RESULTS

### 3D light sheet imaging reveals enhanced brain parenchymal delivery of TfR1-Aducanumab

A single Fab fragment binding to the murine TfR1 was attached to the Fc effector domain of Adu (TfR1-Adu, Fig. 1A). An initial 3D LSFM imaging study was performed to evaluate brain biodistribution of Adu and TfR1-Adu as compared to standard anti-human IgG (Ctrl-hIgG) after single systemic administration in amyloid plaque-bearing mice expressing neuron-specific mutant forms of human APP and PS1 (APP/PS1 mice)^36^ (Fig. 1B). Antibodies were administered intravenously at a relatively high dose (50 nmol/kg) and central spatial distribution of the peripherally administered antibodies was tracked by whole hemibrain labelling of hIgG-positive fluorescent signal (Fig. 1C). 3D reconstructed sagittal/coronal sectional views and fly-through videos reveal the striking differences in brain distribution profiles of Ctrl-hIgG, high-dose Adu and high-dose TfR1-Adu (Fig. 1C-E, Suppl. Movie 1). As expected for a standard IgG, brain accessibility of Ctrl-hIgG was highly restricted. The Ctrl-hIgG signal lined the cerebral ventricles, preferentially accumulating within the choroid plexus, a main component of the blood-cerebrospinal fluid barrier, and fibre tracts covering the ventricular surface (fimbria). In comparison, Ctrl-hIgG signal was considerably weaker in deeper areas of the brain (Fig. 1C-E). Compared to Ctrl-hIgG, a significantly greater Adu signal was detected within the ventricular system and choroid plexus, but not the cerebral cortex and hippocampal region. While both Adu and TfR1-Adu showed higher signal in area postrema, TfR1-Adu signal was substantially lower in the subfornical organ. All other components of the circumventricular organs (CVOs), which are highly fenestrated and vascularized midline structures, showed similar signal of Ctrl-hIgG, Adu and TfR1-Adu. The TfR1-binding shuttle enhanced BBB penetration and brain parenchymal delivery of Adu, as demonstrated by a two-fold increase of TfR1-Adu signal in the cerebral cortex, but not hippocampal region, with a concomitant 6-fold lowering of ventricular signal (Fig. 1C-E, 1F). TfR1-Adu and Adu was detected in a total 245 and 99 brain regions, respectively, with overlapping fluorescent signal in 76 brain regions (Fig. 1G). Terminal plasma exposure analysis indicated that circulating levels of Adu and TfR1-Adu were markedly lower compared to Ctrl-hIgG. Plasma TfR1-Adu levels after single intravenous infusion were 8-fold and 4-fold lower compared to Ctrl-hIgG and Adu, respectively (Fig. 1H).

**Figure 1.**
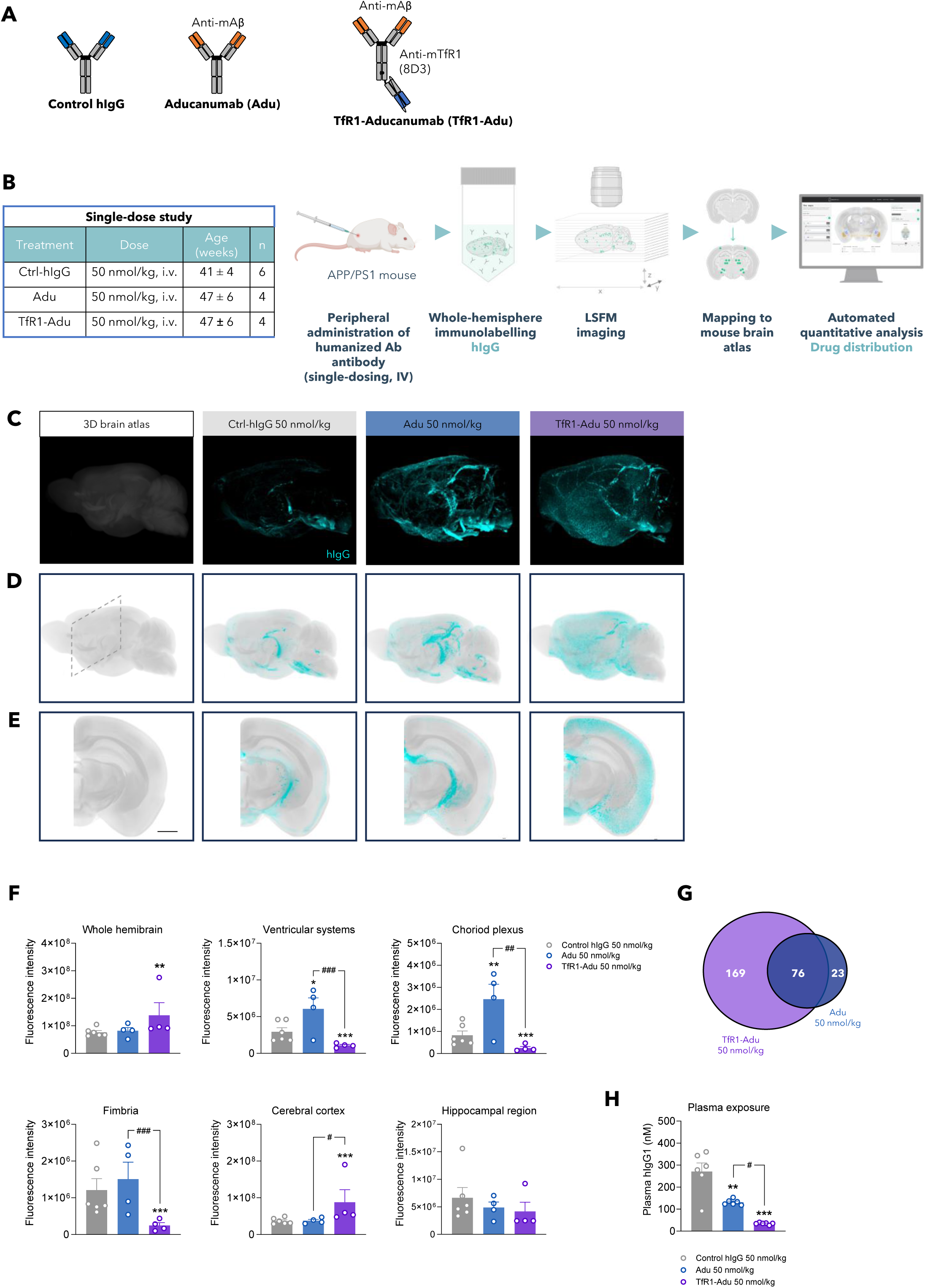
3D LSFM imaging reveals enhanced CNS delivery of TfR1-Aducanumab after single intravenous infusion in APP/PS1 mice. **(A)** Design of humanized IgG antibodies. Schematic illustration of control human IgG (Ctrl-hIgG), Aducanumab (Adu) and Aducanumab fused with anti-mouse transferrin receptor (TfR1-Adu). **(B)** Treatment groups and 3D quantitative LSFM imaging pipeline. Animals were terminated 48h after receiving a single intravenous infusion of Ctrl hIgG (50 nmol/kg, n=6), Adu (50 nmol/kg, n=4) or TfR1-Adu (50 nmol/kg, n=4). Whole hemibrains were stained with anti-human IgG to visualize, map and quantify brain distribution of therapeutic Aβ antibodies after peripheral administration. **(C)** Representative LSFM raw images of anti-human hIgG stained whole hemibrains (sagittal view). **(D)** Virtual 3D samples depicting group-averaged fluorescent signal intensity (sagittal view). Dotted frame indicates rostro-caudal level of virtual 2D coronal sections shown in panel E (scale bar, 30 µm). **(E)** Group-averaged virtual 2D coronal sections sampled at the hippocampal level. **(E)** Accumulated fluorescence signal intensity (arbitrary units) in the whole hemibrain, ventricular systems, choroid plexus, fimbria, cerebral cortex and hippocampal region. **p<0.01, ***P<0.001 vs. Control-hIgG; ^#^p<0.05, ^##^p<0.01 (Dunnett’s test negative binomial generalised linear model). **(F)** Venn diagram depicting distinct and overlapping brain regions with significantly increased TfR1-Adu and Adu signal, as compared to Ctrl-hIgG (FDR<0.05). **(H)** Plasma exposure of Control-hIgG, Adu and TfR1-Adu, measured 48h after single intravenous infusion. **p<0.01, ***p<0.001 vs. Control-hIgG (one-way ANOVA with Dunnett’s post-hoc test).

### Improved brain parenchymal delivery of chronic low-dose TfR1-Aducanumab administration

Because the TfR1 shuttle enhanced brain parenchymal delivery of single intravenous high-dose Adu, we asked if improved brain exposure could be achieved using a 5-fold lower dose of TfR1-Adu in a chronic treatment regimen. To this end, low-dose TfR1-Adu (10 nmol/kg) or low/high-dose Adu (10 nmol/kg, 50 nmol/kg) was administered intraperitoneally once weekly for 12 weeks in APP/PS1 mice, whereafter the left hemibrain was co-stained with anti-hIgG and anti-hAβ (Fig. 2A). Fly-through videos illustrate the brain distribution profiles of Ctrl-hIgG, Adu and TfR1-Adu upon chronic dosing (Suppl. Movie 2). Brain accessibility of low-dose Adu was only marginally greater than observed for Ctrl-hIgG. A five-fold higher dose of Adu improved brain delivery, nevertheless, for both Adu doses a clear IgG signal remained associated with the ventricular systems/choroid plexus (Fig. 2B-C). A greater anatomical overlap in distribution signal was observed for high-dose Adu vs. low-dose TfR1-Adu (Fig. 2D). Nevertheless, low-dose TfR1-Adu showed further enhanced brain delivery compared to both doses of Adu (Fig. 2B-C). Accordingly, TfR1-Adu demonstrated a total hemibrain signal that was 5.1- and 1.4-fold higher compared to low-dose and high-dose Adu, respectively (Fig. 2D). Both Aβ-targeted antibodies accumulated in the cerebral cortex and hippocampal region exhibiting substantial plaque load (Fig. 4). A TfR1-Adu signal was nearly undetectable the ventricular systems/choroid plexus, further supporting improved BBB penetration (Fig. 2D). It is noteworthy that Adu signal was clearly and consistently associated with cerebral surface arteries which was not observed for TfR1-Adu (Fig. 2E, S1). Most striking differences in antibody distribution profiles at the subregional level were observed within the cerebral cortex, where low-dose TfR1-Adu consistently exhibited a greater ability to reach all layers (Fig. 2F). While this was exceptionally clear for low-dose TfR1-Adu vs. low-dose Adu, high-dose Adu also demonstrated less signal accumulation within the deepest cortical layers, layers 5-6 (Fig. 2G).

**Figure 2.**
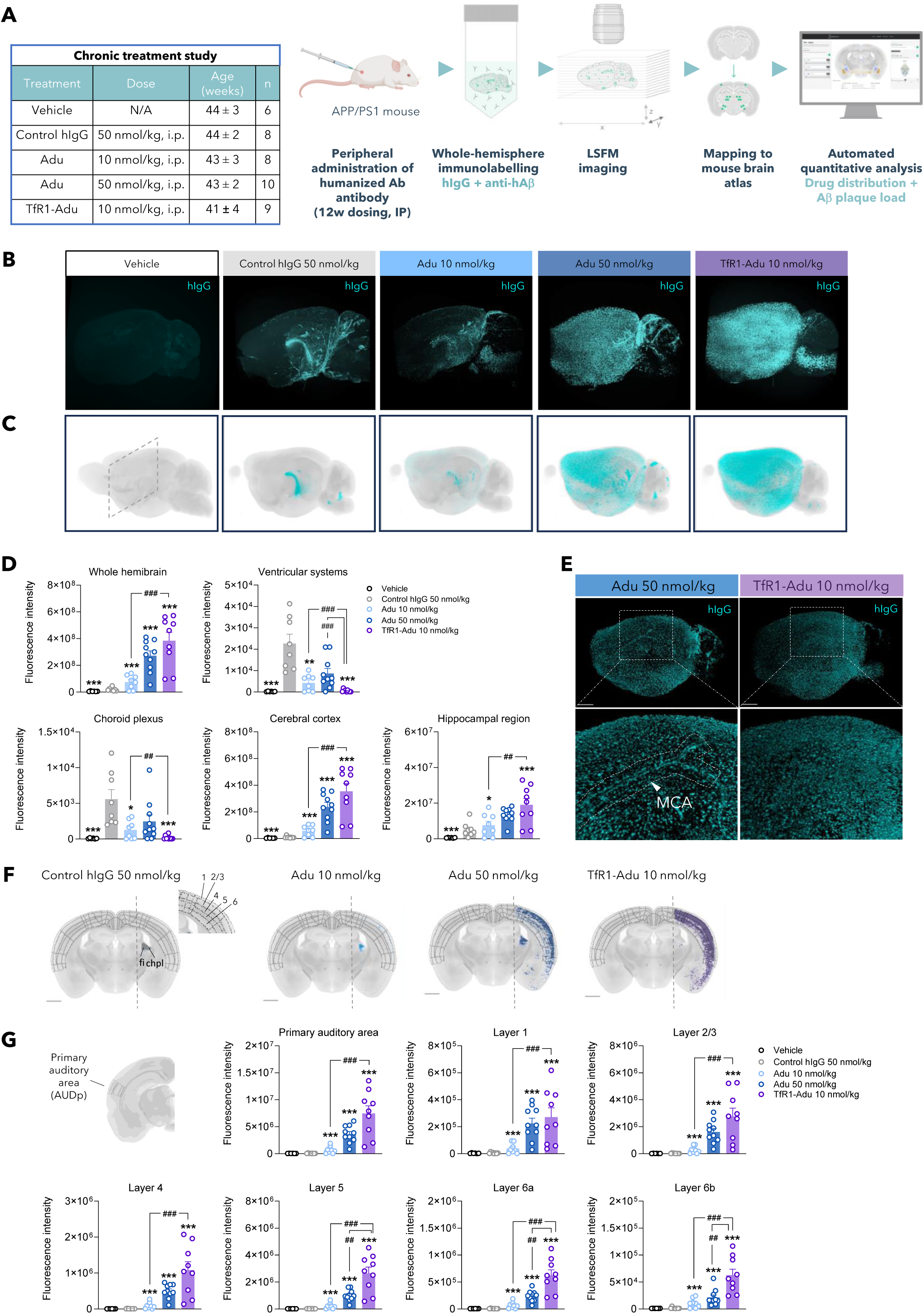
Comparative brain distribution maps of TfR1-Aducanumab and Aducanumab after chronic treatment in APP/PS1 mice. **(A)** Treatment groups and 3D quantitative LSFM imaging pipeline. Animals were terminated 72h after receiving chronic intraperitoneal administration of Ctrl hIgG (n=8), aducanumab (Adu) 10 nmol/kg (n=8), Adu 50 nmol/kg (n=10) or TfR1-Aducanumab (TfR-Adu, 10 nmol/kg (n=9) once weekly for 12 weeks. Whole hemibrain were co-stained with anti-human IgG and anti-human Aβ to simultaneously visualize, map and quantify brain distribution of hIgG signal and Aβ plaque load. **(B)** Representative 3D LSFM raw images of anti-hIgG stained whole hemibrains (sagittal view). **(C)** Virtual 3D samples depicting group-averaged fluorescent signal intensity (sagittal view). **(D)** Accumulated fluorescent signal intensity (arbitrary units) in the whole hemibrain, choroid plexus, cerebral cortex and hippocampus. **p<0.01, ***P<0.001 Accumulated fluorescence signal intensity (arbitrary units) in the whole-hemibrain, choroid plexus, cerebral cortex and hippocampus. *p<0.05, ***P<0.001 vs. Control-hIgG 50 nmol/kg; ^##^p<0.01, ^###^p<0.001 (Dunnett’s test negative binomial generalised linear model). **(E)** Representative LSFM images showing localization of Adu, but not TfR1-Adu, to large surface arteries such as the medial cerebral artery (MCA). **(F)** TfR1-Adu 10 nmol/kg demonstrates deeper brain distribution as compared to Adu (50 nmol/kg). Virtual 2D coronal sections from group-averaged 3D LSFM-imaged brains illustrating differential brain distribution of Adu and TfR-Adu, notably within the cerebral cortex. Cortical layers 1-6 are indicated. **(G)** Accumulated fluorescent signal intensity (arbitrary units) of individual mAbs in the cortical primary auditory area, layers 1-6. fi, fimbria; chpl, chorioid plexus.

### Differentiated brain biodistribution of low-dose Tfr1-Aducanumab vs. unmodified Aducanumab

A subsequent analysis was performed to allow for an unbiased comparison of the global biodistribution profile of each individual antibody (Fig. 3). Brain areas were categorized according to those exhibiting highest average IgG signal of Ctrl-hIgG (cluster 1, 5 regions), TfR1-Adu (cluster 2, 112 regions), and Adu (cluster 3, 15 regions), respectively (Fig. 3A, 3B, Fig. S2). The cluster analysis confirmed limited brain distribution of Ctrl-hIgG, being restricted to the choroid plexus and ventricle-near fiber tracks (fimbria, optic tract) (Fig. 3C). While a Ctrl-hIgG signal was also associated with few discrete thalamic subregions lining the third ventricle (lateral dorsal nucleus of thalamus, subgeniculate nucleus), we cannot exclude if this could potentially be a ‘spill-over’ signal from the ventricle. Most prominent TfR1-Adu signals were detected within the isocortex (36 regions; including the insular cortex and several auditory, somatosensory, visual and motor subdivisions) and subfields of the hippocampal formation (17 regions; including CA1-CA2, dentate gyrus, entorhinal subregions, subiculum and associated extensions) (Fig. 3D). Several other major brain divisions also demonstrated a very clear TfR1-Adu signal, such as the cortical subplate (12 regions; including several amygdalar nuclei, claustrum and endopiriform nucleus), fiber tracks (12 regions; including the anterior commissure, corpus callosum, optic radiation and vestibular nerve), cerebral nuclei (10 regions; including nucleus accumbens, caudoputamen, fundus of striatum and substantia innominate), olfactory areas (7 regions; including nucleus of the lateral olfactory tract and piriform areas) and thalamus (4 regions; including medial geniculate complex and suprageniculate nucleus). Brain areas with dominant Adu signal were largely associated with the thalamus (9 regions; including lateral geniculate complex, dorsal part of the lateral geniculate complex, and reticular nucleus of the thalamus), fiber tracks (6 regions; including genu of corpus callosum and brachium of the superior colliculus), hindbrain (4 regions; including Koelliker-Fuse subnucleus and parabrachial nucleus) and cerebral nuclei (4 regions; including the central amygdalar nucleus lateral/medial part and bed nucleus of the accessory olfactory tract) (Fig. 3E). Based on statistical analyses, low-dose TfR1-Adu IgG signals were significantly greater than Adu in 92 regions (low-dose Adu) and 14 regions (high-dose Adu) (Fig. 3A, S2). Differences in brain regional IgG signal intensities of low-dose TfR1-Adu vs. low-dose/high-dose Adu are summarized in Fig. S3 and S4. As for acute dosing, brain delivery of Tfr1-Adu was inversely correlated with plasma exposure in APP/PS1 mice. TfR1-Adu were largely undetectable in plasma at both time points (Fig. S5). In contrast, dose-dependent increases in Adu plasma levels were observed at dosing week 8 and 12. Taken together, TfR1-Adu showed both region- and subregion-specific enhancement of brain biodistribution.

**Figure 3.**
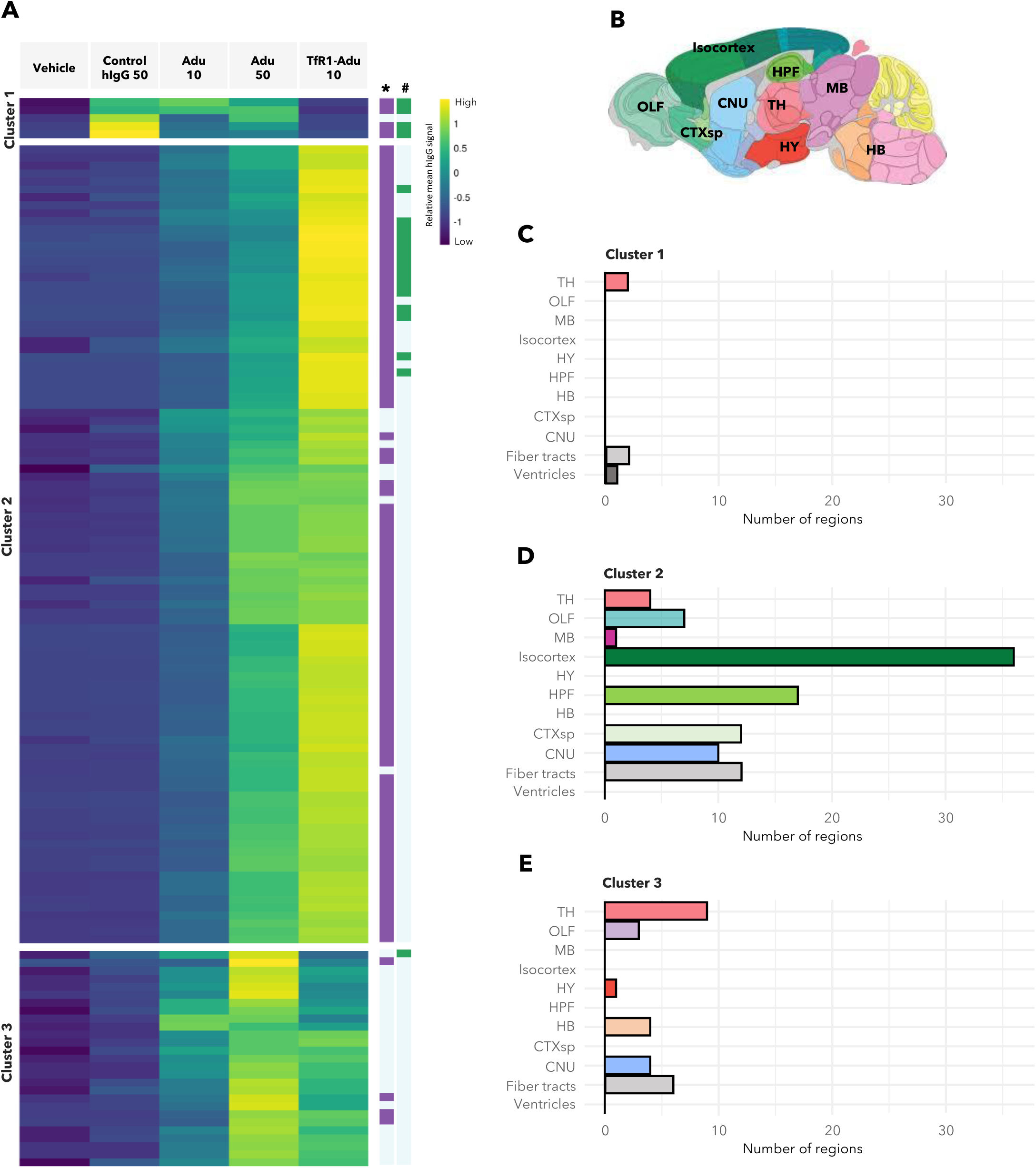
Global brain biodistribution of TfR1-Aducanumab and Aducanumab following chronic treatment in APP/PS1 mice. **(A)** Heatmap of a total of 132 brain regions with increased fluorescence signal of Aducanumab (Adu, 10 nmol/kg, n=8; 50 nmol/kg, n=10) and TfR1-Aducanumab (TfR1-Adu, 10 nmol/kg, n=9) compared to Control hIgG (50 nmol/kg, n=8). Colors indicate relative mean hIgG signal determined in individual brain regions. The heatmap is segmented into three major clusters, using hierarchical clustering, according to brain regions showing highest signal of Ctrl hIgG (cluster 1, n=5 brain regions), TfR1-Adu (cluster 2, n=100 brain regions), and Adu (cluster 3, n=27 brain regions), respectively. *p<0.05 TfR1-Adu (10 nmol/kg) vs. Adu (10 nmol/kg); ^#^p<0.05 TfR1-Adu (10 nmol/kg) vs. Adu (50 nmol/kg), Dunnett’s test negative binomial generalized linear model. **(B)** Brain anatomical distribution maps of therapeutic antibodies according to major CNS regions, including thalamus (TH), olfactory areas (OLF), midbrain (MB), isocortex, hypothalamus (HY), hippocampal formation (HPF), hindbrain (HB), cortical subplate (CTXsp), cerebral nuclei (CNU), fiber tracts and ventricles. **(C-E)** Major brain divisions represented in the three clusters.

### Comparable plaque clearing efficacy of low-dose TfR1-Aducanumab and high-dose Aducanumab

Quantitative 3D LSFM imaging of brain hemibrains co-immunolabeled for hIgG and hAβ provided a unique opportunity to simultaneously map Aβ-targeting vs. Aβ-clearing efficacy of Adu and TfR1-Adu upon chronic treatment in APP/PS1 mice. APP/PS1 mice demonstrated substantial plaque load at study termination, corresponding to 41-44 weeks of age (group-averaged). The distribution of the Aβ-targeted antibodies closely mirrored the distribution of amyloid plaques, indirectly supporting target engagement (Fig. 4). Fly-through videos further illustrate the overlap between drug and plaque deposition (Suppl. Movie 3). Consistent with previous reports on vascular amyloid deposition in other APP/PS1 transgenic mouse models of AD^36^, we observed a putative perivascular signal for Aβ labelled plaques in the APP/PS1 mice, thus presenting with cerebral amyloid angiopathy (CAA) lesions. Notably, the perivascular Aβ signal showed IgG co-labelling for Adu (Fig. 4B, C), but not for TfR1-Adu (Fig. 4D). This further supports our observation that TfR1-Adu appears to efficiently cross the brain capillary endothelium to prevent or limit accumulation within the vasculature. Next, we performed a region-by-region analysis of amyloid plaque counts in all samples evaluated for global Ctrl-hIgG, Adu and TfR1-Adu biodistribution. TfR1-Adu showed plaque-clearing efficacy which could be demonstrated even at the whole hemibrain level, particularly driven by lowered overall cortical plaque load (Fig. 5A). A relatively larger proportion of brain regions demonstrated significantly reduced plaque load following low-dose TfR1-Adu treatment (14 regions) as compared to low-dose Adu (5 regions), but not high-dose Adu (17 regions) (Fig. 5B-D, 6A). 13 brain regions showed significantly reduced Aβ plaque load after both TfR1-Adu and Adu monotherapy, respectively (Fig. 6A). Top-10 brain regions with pronounced reductions in β-amyloid plaque load after chronic dosing with TfR1-Adu or Adu included several cortical and amygdalar subregions (Fig. S6). A scrutinized analysis indicated no region-wise correlation between IgG signal intensity and relative reductions in plaques counts (Fig. 5B-D, Fig. S7). The apparent dissociation between plaque-lowering efficacy and therapeutic antibody exposure at single-region level suggests that brain regions are not equally responsive to Aβ-targeted Immunotherapeutics despite efficient drug delivery.

**Figure 4.**
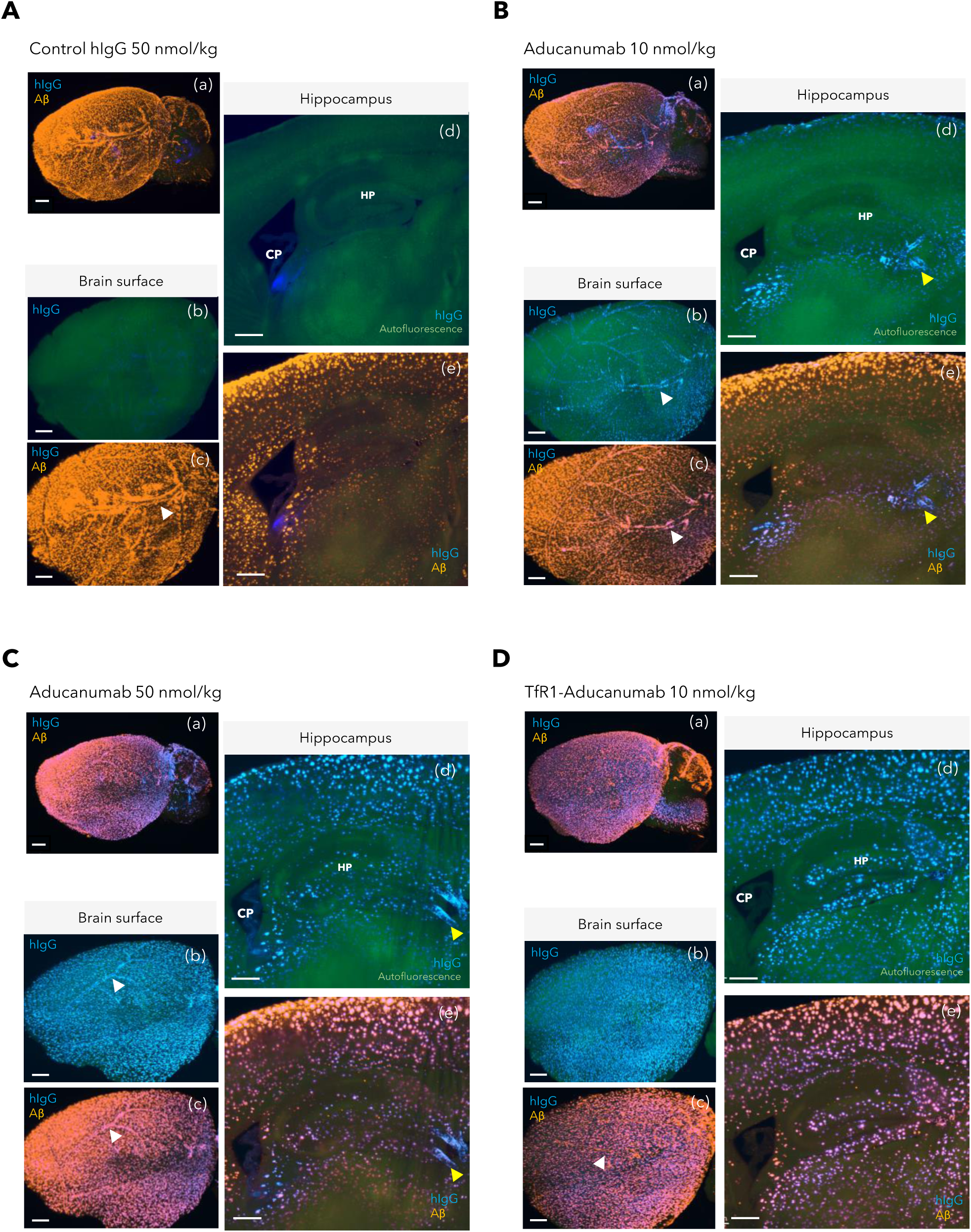
Differential plaque-associated compartmentalization of TfR1-Aducanumab and aducanumab after chronic treatment in APP/PS1 mice. Whereas Aducanumab distributes to both parenchymal and vascular plaques, TfR1-Aducanumab is only localized to parenchymal plaques. Whole hemibrains were co-stained with anti-human IgG and anti-human Aβ to simultaneously visualize, map and quantify brain distribution of therapeutic antibodies and Aβ plaques following treatment with TfR1-Aducanumab or Aducanumab once weekly for 12 weeks. Representative 3D LSFM-imaged brains of mice administered **(A)** control hIgG (50 nmol/kg), **(B)** aducanumab (Adu, 10 nm/kg), **(C)** aducanumab (50 nmol/kg) or **(D)** TfR1-aducanumab (TfR1-Adu, 10 nmol/kg), respectively. *(a)* 3D LSFM sagittal view of whole hemibrain labelled with anti-human IgG (therapeutic antibody distribution, blue color) and anti-human Aβ (amyloid plaques, orange color). *(b)* Close-up view of brain surface distribution of hIgG signal (blue color). *(c)* Combined close-up view of brain surface distribution of hIgG signal and β-amyloid plaques. Scale bar, 700 µm. *(d-e)* 2D virtual sections (100 µm thickness) sampled at the level of the hippocampus (HP) and choroid plexus (CP, lining the lateral ventricle). Scale bar, 200 µm. Arrows indicates vascular-associated deposition of therapeutic antibody and amyloid plaques, respectively.

**Figure 5.**
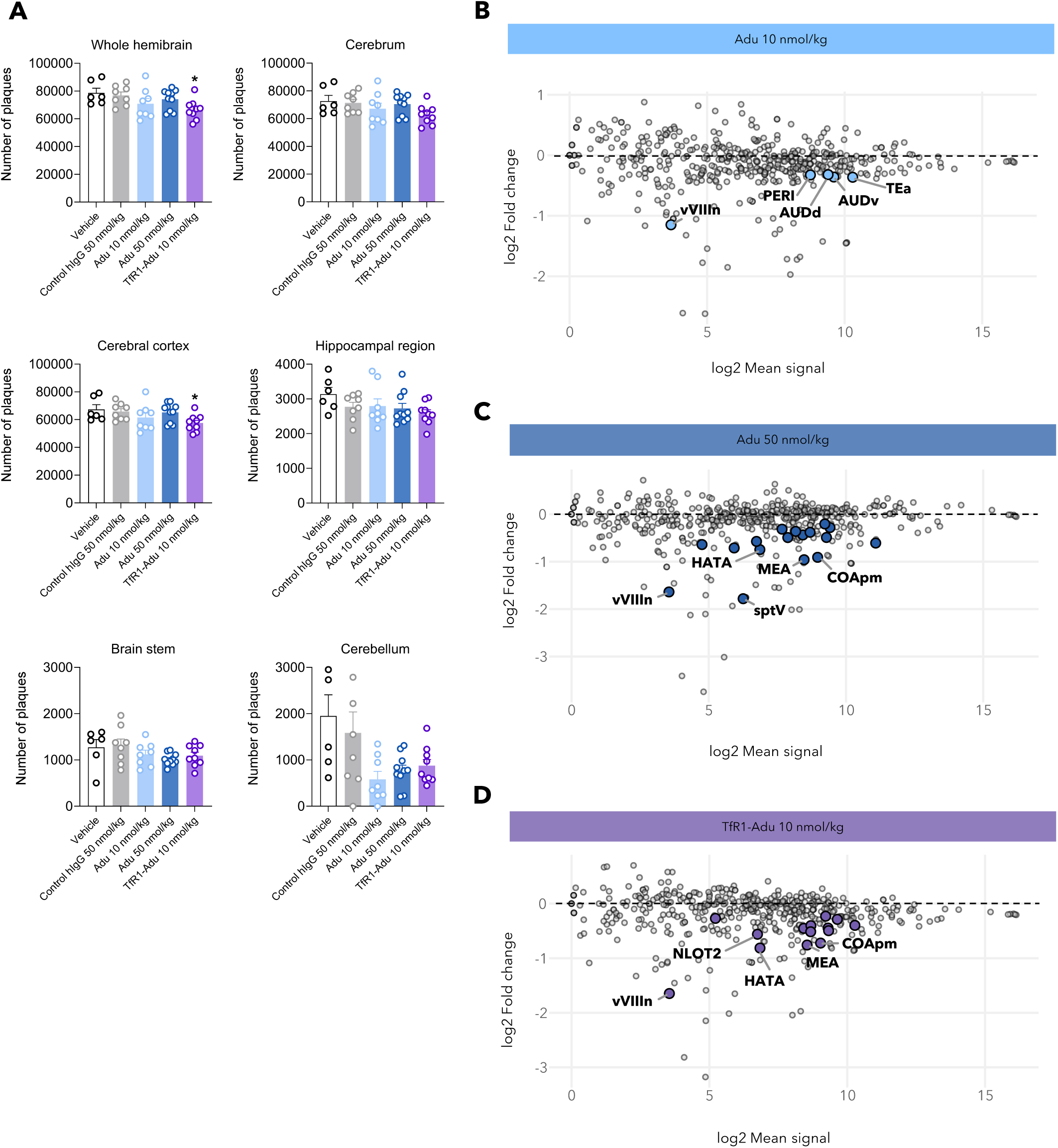
Enhanced Aβ plaque-clearing efficacy of TfR1-Aducanumab. **(A)** Total plaque counts in the whole hemibrain, cerebrum, cerebral cortex, hippocampus, brain stem and cerebellum. **(B-D)** MA-plots displaying log2 mean IgG signal vs. log2-fold change in plaque counts relative to control hIgG (50 nmol/kg) in 840 individual brain regions. Coloured dots indicate regions with significantly reduced plaque counts (p<0.05, Dunnett’s test negative binomial generalized linear model). Top-5 regions with most reduced plaque counts relative to control hIgG are indicated: AUDv (ventral auditory area), AUDd (dorsal auditory area), COApm (cortical amygdala, posterior part, medial zone), HATA (hippocampo-amygdalar transition area), MEA (medial amygdalar nucleus), NLOT2 (nucleus of the lateral olfactory tract, pyramidal layer), PERI (perirhinal area), sptV (spinal tract of the trigeminal nerve), TEa (temporal association areas), vVIIIn (vestibular nerve).

**Figure 6.**
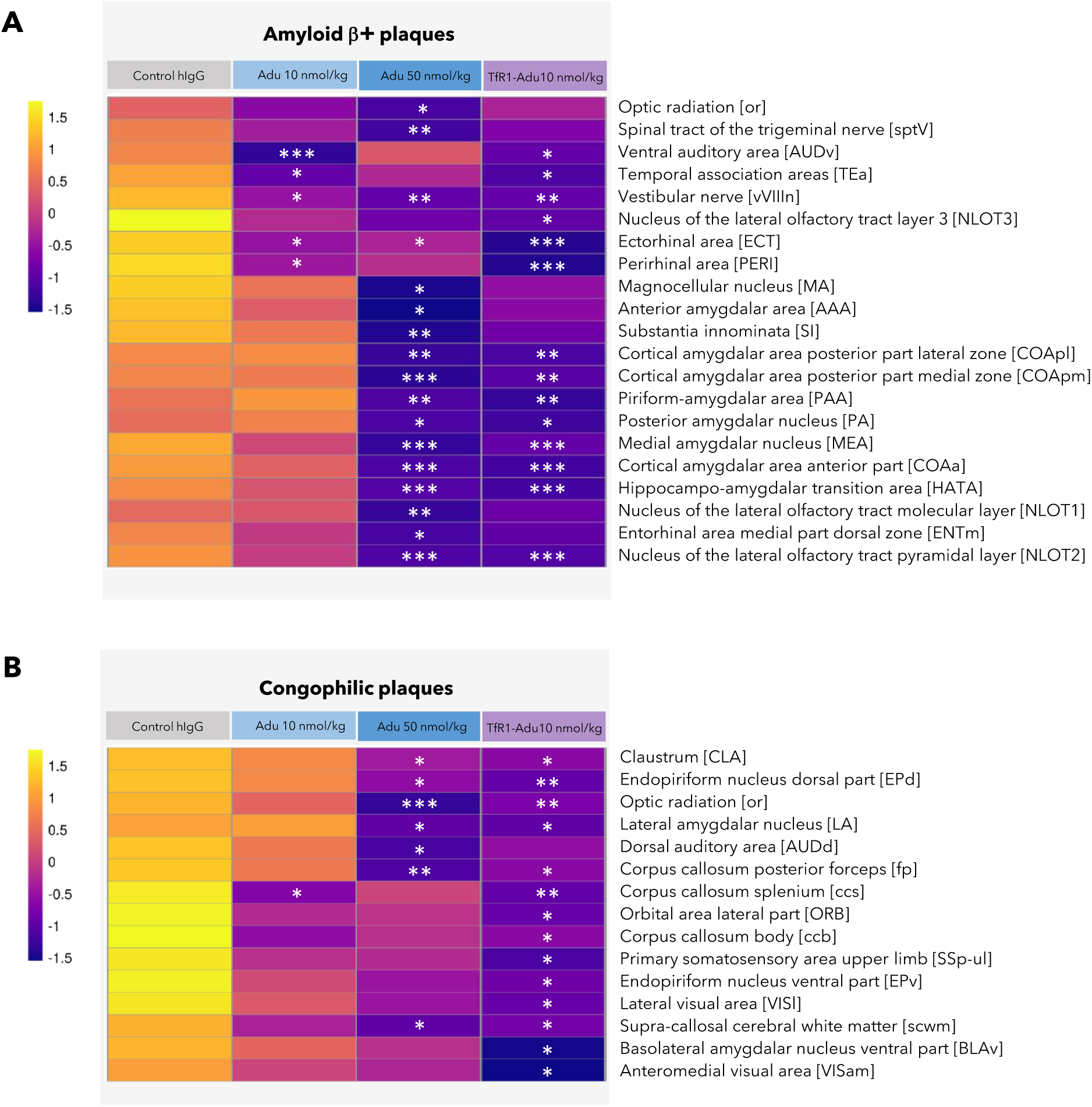
Global comparative analysis of Aβ+ and congophilic plaque burden after chronic dosing with TfR1-Aducanumab or Aducanumab in transgenic APP/PS1 mice. Plaque load was assessed by whole hemibrain labelling with **(A)** anti-human Aβ (Aβ+ plaques, left hemibrain), or **(B)** Congo Red (congophilic plaques, right hemibrain). Brain regions included in the heatmaps showed significant reduction of plaque counts following Aducanumab or TfR1-Aducanumab treatment. *p<0.05, **p<0.01, ***p<0.001 vs. Ctrl hIgG (Dunnett’s test negative binomial generalized linear model).

### TfR1-Adu shows greater capacity to alleviate congophilic plaque burden

There is an increasing appreciation that more mature, compact and fibrillar plaques are more neurotoxic than smaller diffuse plaques^37,38^. We therefore sought to map chronic TfR1-Adu and Adu treatment effects on mature plaques in APP/PS1 mice. To this end, the intact right whole hemibrain was stained with Congo red, a fluorescent dye which only detects dense-core amyloid plaques, *i.e.* senile plaques with a more mature form. Congophilic plaques were detected in all parts of the APP/PS1 mouse brain but were overall at lower counts than Aβ+ plaques (Fig. S7, S8). A greater proportion of brain regions demonstrated significantly reduced congophilic plaque counts following low-dose TfR1-Adu treatment (14 regions) as compared to low-dose Adu (1 region) and high-dose Adu (7 regions) treatment (Fig. 6B). Only optic radiation (also known as the geniculocalcarine tract, the most prominent white matter relay in the visual system) showed significantly reduced Aβ+ and congophilic plaque burden after TfR1-Adu and Adu treatment (Fig. 6A, 6B). In summary, the broader therapeutic effect of TfR1-Adu on mature plaques demonstrates that various plaque populations responded more favourably to TfR1-Adu.

## DISCUSSION

The current quantitative 3D imaging study aimed to characterize the impact of a TfR1 BBB shuttle on brain delivery and efficacy of Adu, a clinically relevant anti-Aβ mAb. We here report superior brain parenchymal distribution and plaque clearing efficacy of TfR1-Adu in a transgenic mouse model of AD. Taken together, our data lends support for TfR1-targeting being a valid strategy for enhancing brain uptake and therapeutic efficacy of Aβ-targeted immunotherapeutics.

High-throughput quantitative 3D LSFM imaging coupled with computational analysis of iDISCO-cleared whole hemibrains enabled detailed simultaneous assessment of brain delivery, biodistribution and plaque clearing efficacy of peripherally administered TfR1-Adu and unmodified Adu in Aβ plaque-bearing transgenic APP/PS1 mice. Our study establishes that TfR1-Adu reaches deep into the brain tissue and showed persistence of intensity and broad parenchymal signal distribution regardless of relatively fast systemic clearance. Together, this is indicative of enhanced brain uptake, biodistribution and retention of TfR1-Adu compared to unmodified Adu. Interestingly, high signals of Ctrl-hIgG and Adu, but not TfR1-Adu, were detected in cerebrospinal fluid (CSF)-bordering areas, particularly the choroid plexus which is a highly vascularized and TfR1-enriched structure with prominent blood supply^39,40^. It is possible that retention of Ctrl-hIgG and Adu in the choroid plexus was accentuated by FcRn receptor-mediated export, which has been suggested to play an important role in IgG antibody clearance from the brain parenchyma to the circulation^41,42^. Accumulation of Ctrl-hIgG and Adu in the choroid plexus most likely reflects lack of specialized BBB transport for these antibodies. Restricted brain access of Ctrl-hIgG is consistent with previous studies demonstrating prominent periventricular and perivascular localization of standard IgGs with highly limited parenchymal biodistribution, even at very high doses administered systemically or via intracisternal route^43,44^. This suggests minimal BBB passage of standard IgGs and gain access to superficial brain areas via the blood-to-CSF trafficking route. While standard IgGs specifically localize to barrier cells such as choroid plexus epithelial cells, meningeal fibroblasts and perivascular immune cell populations, TfR-enabled anti-Aβ mAbs distribute to neurons and glial cells via transcytosis through neurovascular endothelial cells^43^. Collectively, we therefore infer that improved brain delivery and widespread parenchymal accumulation of TfR1-Adu is likely governed by efficient BBB passage, possibly augmented by enhanced TfR1-Adu penetration from the CSF.

Unexpectedly, limited brain delivery and biodistribution of unmodified Adu could be overcome with chronic administration of a five-fold higher systemic Adu dose. Our findings contrast the notion that the BBB is nearly impermeable for large biologics, including antibody therapeutics^11–13^. While APP/PS1 mice demonstrated perivascular plaque accumulation, a clear sign of CAA which is an integral component of AD and associated with cerebrovascular dysfunction and microhaemorrhages^45,46^, it should be noted that transgenic APP and APP/PS1 mouse models display intact BBB or minimal BBB disruption^47–50^. This makes it unlikely that increased brain access of unmodified Adu could be explained by compromised BBB integrity. Our data align well with earlier studies in transgenic APP mice which estimated that brain clearance of the Aβ antibodies bapineuzumab and gantenerumab was up to eight times slower than in the circulation^51,52^. This raises the intriguing possibility that anti-Aβ mAbs have a unique brain PK profile due to their high amyloid plaque affinity. 3D LSFM imaging of hIgG-hAβ co-immunolabelled whole hemibrains revealed a consistent overlap between the distribution of TfR1-Adu/Adu and the deposition of hAβ+ plaques in APP/PS1 mice. The finding that amyloid plaque distribution likely dictates brain biodistribution of these Aβ-directed antibodies is an important insight. By showing this overlap, our study suggests that anti-Aβ mAbs are preferentially binding to areas where amyloid plaques are present, and that long-lasting plaque engagement is a prerequisite for brain parenchymal retention of these antibodies.

Enhanced brain delivery and parenchymal biodistribution of TfR1-Adu translated to improved plaque clearing efficacy in APP/PS1 mice as compared to dose-matched Adu. Plaque segmentation, mapping and quantification revealed that low-dose TfR1-Adu reduced amyloid plaque populations to a greater extent than observed for both low-dose Adu (Aβ+ and congophilic plaques) and high-dose Adu (congophilic plaques). Similar to our quantitative LSFM 3D imaging data, previous 2D histological studies in other transgenic AD mouse models have reported relatively modest improvements in plaque counts following treatment with Adu^2,53–55^. Plaque clearance afforded by TfR1-Adu likely involves glial cell-mediated phagocytosis and degradation through binding to the Fc region, as previously suggested for unmodified Adu and other anti-Aβ mAbs^2,55^. Correspondingly, a TfR-conjugated Aβ mAb has recently been reported to engage microglia and induce microglial phagocytosis of plaques in AD mice harboring the hTfR apical domain (5xFAD; TfR^mu/hu^ knock-in mice)^56^. There was a clear representation of cortical areas with reduced plaque burden after long-term low-dose TfR1-Adu treatment, being consistent with cortex-wide distribution of TfR1-Adu. As for brain parenchymal delivery, a weaker therapeutic effect of unmodified Adu could be overcome by administering a 5-fold higher dose.

It should be emphasized that TfR1-Adu lowered plaque levels in only a subset of brain regions. The regionally restricted plaque clearing efficacy of TfR1-Adu contrasts the clear evidence of brain-wide delivery of TfR1-Adu. As we did not observe a correlation between region-wise IgG signal intensity and relative reductions in plaques counts, we were unable to determine if there exists a lower threshold level of parenchymal anti-Aβ antibody exposure to facilitate plaque clearance. It should be emphasized that chronic TfR1-Adu therapy was evaluated in 7-8-month-old APP/PS1 mice, which at this age have been reported to carry extensive amyloid lesions dominated by large-size amyloid deposits surrounded by diffuse (small, immature) plaques^36^. As benefits on congophilic plaques were more limited than for Aβ+ plaques, we speculate that TfR1-Adu and Adu might preferentially deplete newly formed plaques while being less effective against existing mature plaques in older APP/PS1 mice. In support of this notion, repeated administration of hAβ mAbs, including Adu, more efficiently clears and/or prevents small plaques over large fibrillar plaques in transgenic AD mouse models^55,56^. Future studies must therefore aim to clarify if plaque depletion afforded by TfR1-Adu treatment might be even more pronounced at earlier stages of amyloidosis.

As opposed to unmodified Adu, plasma TfR1-Adu levels were almost undetectable after chronic dosing suggesting that the TfR1 moiety altered peripheral pharmacokinetic properties of Adu. TfR1 expression is not unique to the BBB, and TfR1 binding with subsequent internalization in other tissues could constitute a peripheral sink for TfR1-Adu, a mechanism referred to as target-mediated drug disposition and degradation^57,58^. In accordance, fast blood clearance and high uptake in the liver, spleen, kidney and blood cells has been observed for TfR1-Adu immuno-PET ligands and other bispecific TfR1-Aβ mAbs in transgenic APP and APP/PS1 mice, presumably explained by TfR1 interaction with the mouse 8D3 moiety^59–62^. Anti-drug antibody (ADA) formation directed towards the TfR1 or Adu moiety may potentially have contributed to enhanced TfR1-Adu clearance. Previous studies have demonstrated that hIgG administration in mice promotes murine anti-hIgG responses, likely due to foreign epitopes present in hIgG proteins, which can develop within days to several weeks of administration and lead to enhanced clearance of the hIGg^63,64^. Additionally, ADA formation has been reported following treatment with a TfR-hAβ mAb (RmAb158-scFv8D3) in wildtype mice, targeting the bispecific mAb for hepatic internalization and catabolic clearance^61,62^. It should be noted considered that immunogenicity was specifically ascribed to the reformatted TfR1-binding 8D3 moiety (scFv8D3), making it unresolved if ADA responses could also develop towards the unmodified 8D3 Fab moiety in TfR1-Adu.

Clinical trials have identified vascular-associated neurological complications, termed amyloid-related imaging abnormalities (ARIA) in the form of MRI-determined oedema/effusions or haemorrhagic lesions, as the major safety concern for Aβ immunotherapies^1,3,7,65^. ARIA liability of Aβ immunotherapies significantly challenges patient selection and restricts implementation of this promising drug class to the broader AD patient population^66^, making it pertinent to develop more both safer and more effective Aβ-directed therapies. Although the neurovascular pathological mechanisms are poorly understood, co-existing CAA is considered the major risk factor for ARIA adverse events with current Aβ immunotherapies^67^. Preclinical evidence suggests that CAA-Aβ antibody immune complex formation triggers inflammatory perivascular immune cell responses which exacerbate CAA-related vascular leakage and microhemorrhages^68^. It is therefore noteworthy that our study in APP/PS1 mice demonstrated that anti-TfR1 conjugation mitigated arterial labelling of Adu, inviting the possibility that TfR1-enabled Aβ immunotherapies could protect against ARIA events while alleviating amyloidosis. Correspondingly, the incidence of ARIA-like lesions and vascular inflammation was significantly reduced with a TfR-Aβ antibody in a mouse model of AD-CAA, being ascribed to less interaction with arterial amyloid^56^. In support of clinical relevance, recent interim data from a phase-1b/2a clinical trial data with trontinemab, a TfR1-enabled Aβ antibody, indicated rapid depletion of amyloid plaques while provoking few cases of mild ARIA in patients with prodromal or mild to moderate AD (#NCT04639050, ClinicalTrials.gov)^69^.

The study has some limitations. The study employed both female and male APP/PS1 mice at different ages and a presumed varying degree of disease severity. A more uniform distribution of plaque burden at baseline could potentially reduce cohort variability and promote further resolution of TfR1-Adu vs. Adu brain delivery and efficacy. Whilst Adu and the polyclonal anti-hAβ antibody applied for LSFM imaging do not bind the same hAβ epitope, we cannot exclude the possibility that Adu-coating might reduce LSFM anti-hAβ antibody affinity leading to an overestimation of the plaque-clearing efficacy of TfR1-Adu and Adu, particularly in plaque-dense regions. Also, the study did not assess potential formation of neutralizing ADAs which could be a contributing mechanism in the systemic clearance of TfR1-Adu. In addition, further studies should aim to establish if lack of arterial labelling of TfR-1-Adu could potentially translate to reduced risk of ARIA-like lesions.

In summary, we report that TfR1 shuttle-enhanced brain parenchymal delivery of Adu allowed for significantly reducing dose levels of Adu without compromising plaque-clearing efficacy in APP/PS1 mice. An additionally important brain distribution feature was the absence of cerebral perivascular labelling of TfR1-Adu. Because Aβ antibody engagement with vascular amyloid deposits is strongly associated with ARIA which has emerged as a major complication of Aβ immunotherapies^1–3,7^, our findings could have important implications for the development of more efficacious and safe immunotherapies for AD.

## METHODS

### Antibody and TfR1 BBB-shuttle synthesis

Antibodies were produced and provided by H. Lundbeck A/S. Adu preferentially targets Aβ plaques but not monomers^2^. The mono- and bispecific antibodies were recombinantly produced as human IgG1/κ (m17.1) using pTT5 vector and transient expression in HEK293 6E cells. The knob-in-hole technology was utilized to engineer the bispecific antibody, TfR1 (8D3)-Aducanumab (TfR1-Adu). The anti-TfR1 (8D3) Fab moiety was genetically fused to the C-terminus of the Fc with the knob mutation (T366W) to target the murine TfR1 and facilitate shuttling across the BBB^70^. The “hole” mutations were following T366S, L368A, Y407V, and Y349C in the other arm. Briefly, expression of antibodies was performed by transfecting HEK293 6E cells with appropriate plasmid using PEIpro as the transfection reagent. The cells were harvested 6 days after transfection. The antibodies were purified using HiTrap protein G, washed in PBS and eluted using 20 mM Acetate buffer (pH 3.5). Elution fractions were immediately buffer exchanged into PBS buffer (pH 7.2) by dialysis. This was followed by a second polish step by SEC. Purity was confirmed using SEC-HPLC, SDS-PAGE and MS. For all antibodies, the endotoxin level was < 5 EU/mg. Quality control and mTfR1 binding of TfR1-Adu has been reported previously^60^.

### Animals

C57BL/6J-Tg(Thy1-APPsw-Thy1-PSEN1*L166P)21 mice were used in all studies. The mice were licensed from Dr. Peter Koesler and breed at Charles River Laboratories GmbH, Germany. The APP/PS1 mice carry human transgenes for amyloid-beta precursor protein (APP, Swedish mutation) and presenilin-1 (PS1, L166P mutation), both under the control of the Thy1 promoter. In these mice, expression of the human APP transgene is approximately 3-fold higher than endogenous murine APP, with human Aβ42 being preferentially generated over Aβ40, but both isoforms increase with age^36^. All animals had *ad libitum* access to water and chow (Brogaarden, Lynge, Denmark) in a controlled environment (light/dark cycle maintained at 12 h; room temperature 21 ± 2 °C, relative humidity 55 ± 5%). The animal experiments were performed in accordance with the European Communities Council Directive no. 86/609, the directives of the Danish National Committee on Animal Research Ethics, and Danish legislation on experimental animals (license no. 2014-15-0201-00339).

### Drug treatment

LSFM-based CNS mapping of therapeutic Aβ antibody distribution was determined after single- and chronic dosing, respectively, in APP/PS1 mice. Mixed genders were used in both studies. In the single-dose study, APP/PS1 mice received an intravenous bolus injection in the tail (50 nmol/kg; 5 ml/kg) of standard human IgG (Ctrl-hIgG, n=6), Aducanumab (Adu, n=4), or TfR1-Aducanumab (TfR1-Adu, n=4). Animals were returned to their home cage and sacrificed 48h after dosing. In a subsequent chronic dosing study, APP/PS1 mice were randomized and stratified according to age to ensure comparable age-ranges in treatment groups. Mice received an intraperitoneal injection (5 ml/kg) of either Ctrl-IgG (50 nmol/kg, n=6), Adu (10 nmol/kg, n=8; 50 nmol/kg, n=10), or TfR1-Adu (10 nmol/kg, n=9) once weekly for 12 weeks. Animals were returned to their home cage and sacrificed 72h after the last dose.

### Tissue sampling

Mice were anesthetized with Avertin (tribromoethanol, 250 mg/kg i.p.). A blood sample was collected from the heart, and plasma was isolated by centrifugation for further analysis of hIgG levels by LC-MS. Mice were transcardially perfused with chilled heparinized PBS followed by 10% neutral-buffered formalin brains were collected and post-fixed overnight before being transferred to PBS with 0.1% sodium azide.

### Whole hemibrain staining and clearing

Hemibrains were washed in phosphate-buffered saline (PBS, pH 7.4) for 3 × 30 min at room temperature and dehydrated in a methanol-H_2_O gradient (20-40-60-80%-100% methanol, each step 1 hour at room temperature). Samples were washed in 100% methanol for 1 hour and incubated overnight in 66% dichloromethane (DCM)/33% methanol at room temperature. Samples were washed twice in 100% methanol for 30 minutes, cooled down to 4°C in 1 hour and bleached in chilled fresh 5% H_2_O_2_ in methanol overnight at 4°C. The samples were subsequently rehydrated in a methanol-PBS series (80-60-40-20%, with 0.2% Triton X-100, each step 1 hour at room temperature) and washed in PBS with 0.2% Triton X-100 for 2×1 hour at room temperature. Samples were incubated in permeabilization solution at 37°C for 3 days. Blocking was carried out in blocking solution at 37°C for 2 days. For combined detection of therapeutic antibody and amyloid plaque distribution, samples were co-incubated with donkey Alexa fluor 790 anti-human IgG (1:400, #709-655-149, polyclonal, Jackson ImmunoResearch, West Grove, PA) and rabbit anti-human Aβ (1:100; #18584, polyclonal, IBL, Minneapolis, MN) in PBS with 0.2% Triton X-100/3% donkey serum/5% DMSO at 37°C for 14 days, followed by a series of washes in PBS with 0.2% Triton X-100 (1×10 minutes, 1×20 minutes, 1×30 minutes, 1×1 hour, 1× 2 hours and 1× 3 days). Samples were then incubated with secondary antibody Cy5 AffiniPure anti-rabbit IgG (1:1000, cat no. 711-175-152, polyclonal, Jackson ImmunoResearch) in antibody buffer for seven days at 37°C. For detection of congophilic plaques, brains were blocked and permeabilized as above, followed by incubation with Congo Red dye (Merck, BioXtra, C6277) in PBS with 0.2% Triton X-100/3% donkey serum/5% DMSO at 37°C for 7 days. This was followed by a series of washes in PBS/0.2% Tween-20 (10 min, 20 min, 30 min, 1 hr, 2 hr, 24 hr). Finally, whole-brain clearing was performed according to the protocol by Renier et al.^71^, however, with slight modifications. In brief, samples were dehydrated in a graded methanol series (20-40-60-80-100% in H_2_O, each step 1 hour at room temperature), and incubated overnight in 100% methanol at room temperature. Samples were then transferred to 66% DCM/33% methanol and incubated for 3 hours at room temperature with shaking. For clearing, tissues were transferred to 100% DCM for 2 × 15 min and stored in dibenzyl ether (DBE) until scanning by LSFM.

### 3D light-sheet fluorescence microscopy

Hemibrains were imaged using Lavision light-sheet ultramicroscope II (Miltenyi Biotec GmbH, Bergisch Gladbach, Germany) with Zyla 4.2PCL10 sCMOS camera (Andor Technology, Belfast, UK), SuperK EXTREME supercontinuum white-light laser EXR-15 (NKT Photonics, Birkerød, Denmark) and MV PLAPO 2×C (Olympus, Tokyo, Japan) objective. Samples were mounted in a silicone-casted sample holder (dorsal side up) and imaged in a DBE filled chamber. ImSpector microscope controller software (v7) was used (Miltenyi Biotec GmbH, Bergisch Gladbach, Germany). Horizontal images were acquired at 1.2 × total magnification and 10 μ m z-stack intervals. Horizontal focusing was captured in nine planes. Autofluorescence images were captured at 560 ± 40 nm (excitation) and 650 ± 50 nm (emission) wavelength (80% laser power in ImSpector software, 100% NKT laser). Amyloid beta signal was imaged at 650 ± 25 nm excitation wavelength and 671 ± 55 nm emission wavelength (100% laser power in software, 100% NKT laser). Human IgG signal was imaged at 785 ± 25 nm excitation wavelength and 845 ± 55 nm emission wavelength (100% laser power in software, 100% NKT laser). For brains stained with Congo red, autofluorescence images were captured at 490 ± 40 nm (excitation) and 525 ± 50 nm (emission) wavelength (80% laser power in ImSpector software, 100% NKT laser), while Congo red signal was captured at 560 ± 40 nm excitation and 650 ± 50 nm emission wavelength (100% laser power in software, 100% NKT laser). To minimize experimental variation, standardized procedures were used for processing of all samples. Hence, samples were cleared simultaneously, oriented identically in the microscope and scanned using the same scanning settings. Light sheets were aligned and calibrated once daily and randomized scanning order between brains from different study groups was applied.

### Image analysis

Region delineation of the whole-brain samples was obtained by atlas segmentation. Atlas annotations were obtained by alignment to a digital LSFM-based mouse brain atlas^72^. The atlas utilizes the common coordinate framework version 3 (CCFv3) developed by the Allen’s Institute of Brain Science^73^ and uses the same nomenclature. Registration between the atlas and the individual samples was performed by a global affine alignment followed by a local multi-resolution B-spline-based alignment, using the Elastix image registration toolbox^74^. The registration was based on the autofluorescence image volumes, which were pre-processed by down sampling to 25 μm isotropic resolution and contrast enhancement by contrast limited adaptive histogram equalization (CLAHE). Spectral unmixing was performed to suppress the contribution of tissue autofluorescence in the acquired specific channel of the full brain samples, allowing for the quantification of the distribution of the human antibodies. Image analysis was conducted in Python. The estimated autofluorescence contribution in the specific channel was calculated and removed based on ratios of voxel intensities between selected voxels in the non-specific channel and the corresponding voxels in the specific channel. An adapted ClearMap^75^ routine was employed to obtain plaque aggregate counts. The routine comprised background subtraction and difference of Gaussians to enhance the plaque morphology, maximum filtering to obtain the plaque centroids used in seeded watershed, followed by plaque aggregate size filtering, which permitted only aggregates within the range of 50 to 500 voxels. Upon atlas registration and signal unmixing, group averages and standard deviations could be calculated at each voxel location, which further allowed p-value calculations compared to the Control-hIgG group. All images and video material were generated using Imaris x64 software v. 10.2.0 (Oxford Instruments, Abingdon, Oxfordshire, England). MA plot and heatmap visualisations were made in R version 3.6.3 using packages ggplot2 and pheatmap, respectively. For biodistribution heatmaps, regions were subject to hierarchical clustering using the hclust function in R.

### Statistics

Statistical analysis was performed following the approach outlined by Perens et al.^72^. To prevent signal redundancy, only regions from the lowest level of the atlas hierarchy were selected. To further reduce the number of regions, some were merged one level up, resulting in a total of 438 regions. For determining the difference across groups, a generalized linear model was fitted to the signal observed in each brain region in every animal group. A negative binomial generalized linear model provided a suitable fit to our data. For each generalized linear model, a Dunnett’s test was performed. Statistical analysis of the data was performed using R statistics library. Further, all significantly regulated brain regions underwent a two-step manual validation procedure for checking if the used statistical model fits the data points, the significance of the brain regions is not achieved due to outliers and the raw signal is truly originating from the region. First, the fit of data to the generalized linear model was evaluated. This was done by investigating deviance residuals and checking if the residuals aligned with the assumptions of normality and homoscedasticity. Furthermore, Cook’s distance was calculated for each cell count/ intensity data point in the model as a measure of model influence. Regions where the generalized linear model showed severe violations of the assumptions, or the model contained overly influential data points, were discarded. Secondly, the remaining brain regions were visually studied for possible spillover signal from neighbouring regions. If the signal response in a region seemed to originate from the neighbouring region, e.g. very few counts were observed only in the border areas of the region while the neighbouring areas were exhibiting remarkably high signal, it was declared as not significant.

## DATA AVAILABILITY

Data are available upon reasonable request.

## ACKNOWLEDGEMENTS

N/A

## AUTHOR CONTRIBUTIONS

H.H.H., S.V, A.J. and J.H.S. conceived the study, designed experiments, interpreted data, co-wrote the manuscript, and edited the manuscript. M.R.V. performed experiments, analyzed and interpreted data, co-wrote and edited the manuscript. C.S.J. designed and performed experiments. F.W. performed experiments and edited the manuscript. E.A., J.L.S., M.R.M. and C.G.S. analyzed data and edited the manuscript. F.S. analyzed data, interpreted data and edited the manuscript.

## ADDITIONAL INFORMATION

The online version contains supplementary video material (DOI 10.6084/m9.figshare.28253876; https://figshare.com/s/ab22f348a615e13ac915).

---

**Figure S1.**
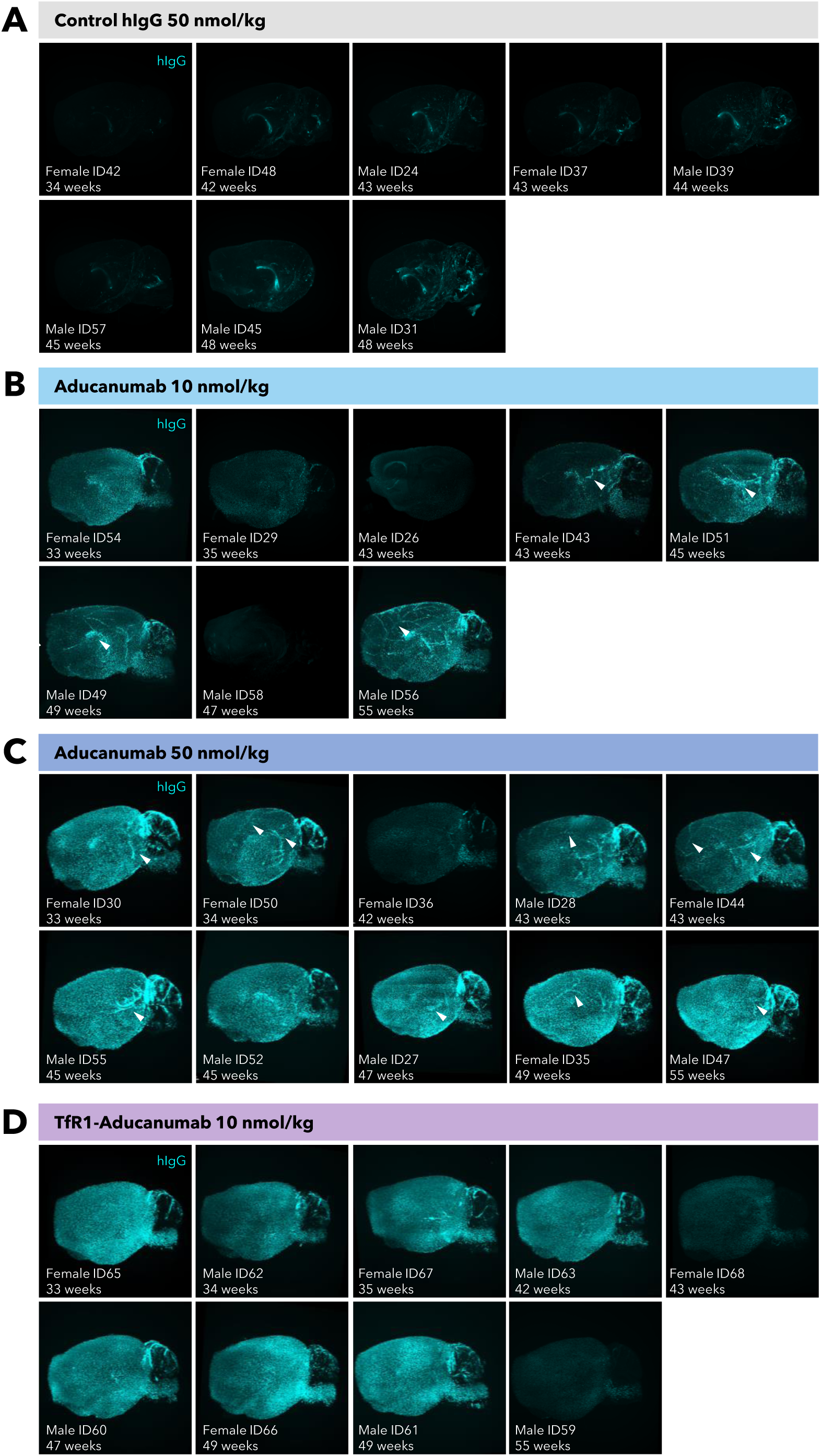
Brain biodistribution of Control hIgG, Aducanumab and TfR1-Aducanumab after chronic dosing in APP/PS1 mice. 3D sagittal projections of LSFM-imaged brains from transgenic APP/PS mice receiving chronic treatment (intraperitoneal administration once weekly for 12 weeks) with **(A)** Control hIgG (50 nmol/kg, n=8), **(B)** Low-dose Aducanumab (10 nmol/kg, n=8), **(C)** High-dose Aducanumab (50 nmol/kg, n=10), **(D)** Low-dose TfR1-Aducanumab (10 nmol/kg, n=9). The left hemibrain was stained with anti-human IgG to determine central distribution of the therapeutic antibodies. Aducanumab, but not TfR1-aducanumab, demonstrates (peri)vascular-associated labelling (white arrows). Gender and age at termination is indicated for each animal.

**Figure S2.**
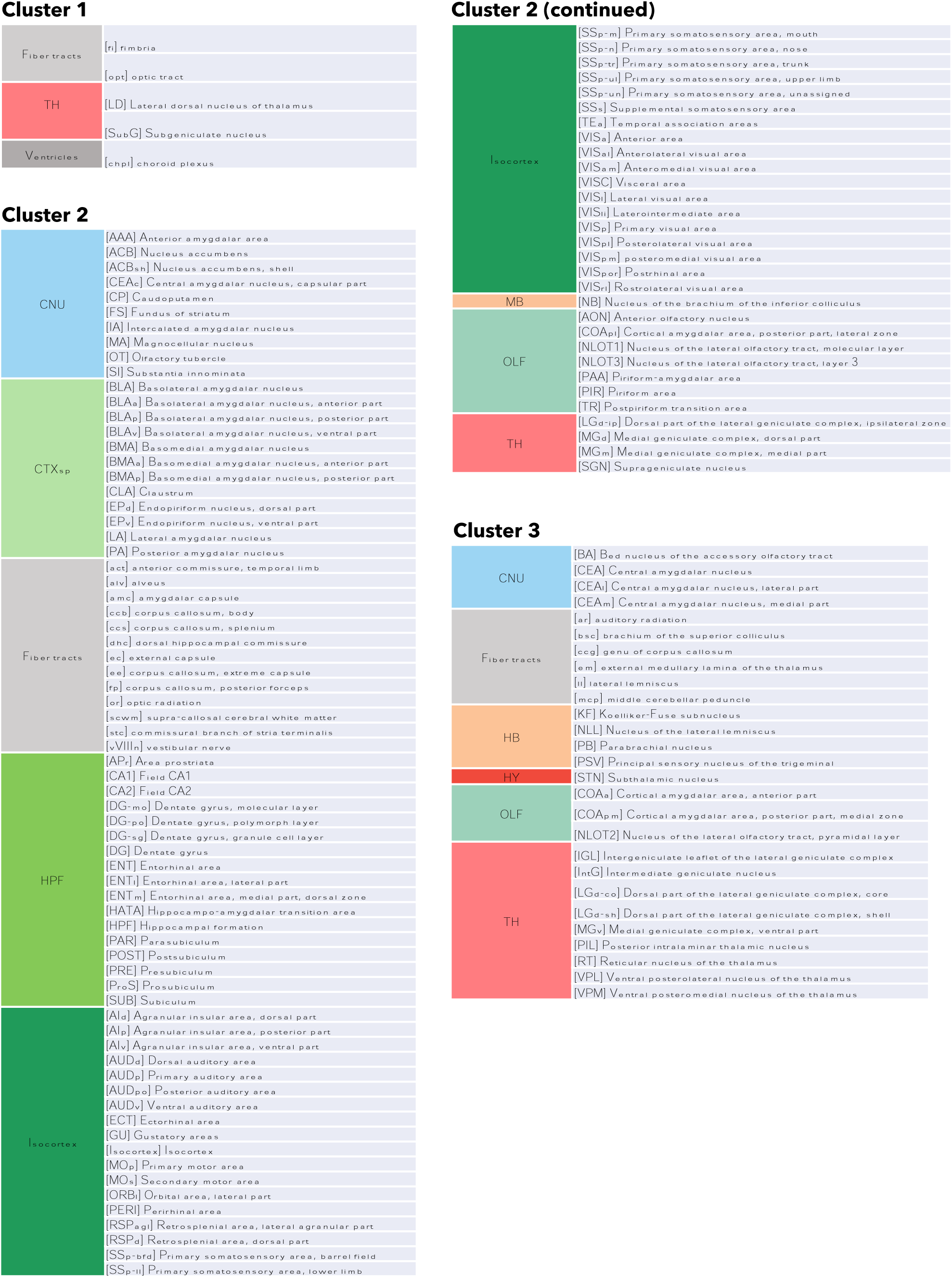
Major brain divisions and associated regions represented in the three clusters of therapeutic antibody distribution. *Abbreviations:* CNU, cerebral nuclei; CTXsp, cortical subplate; HB, hindbrain; HPF, hippocampal formation; HY, hypothalamus; MB, midbrain; OLF, olfactory areas;, TH, thalamus.

**Figure S3.**
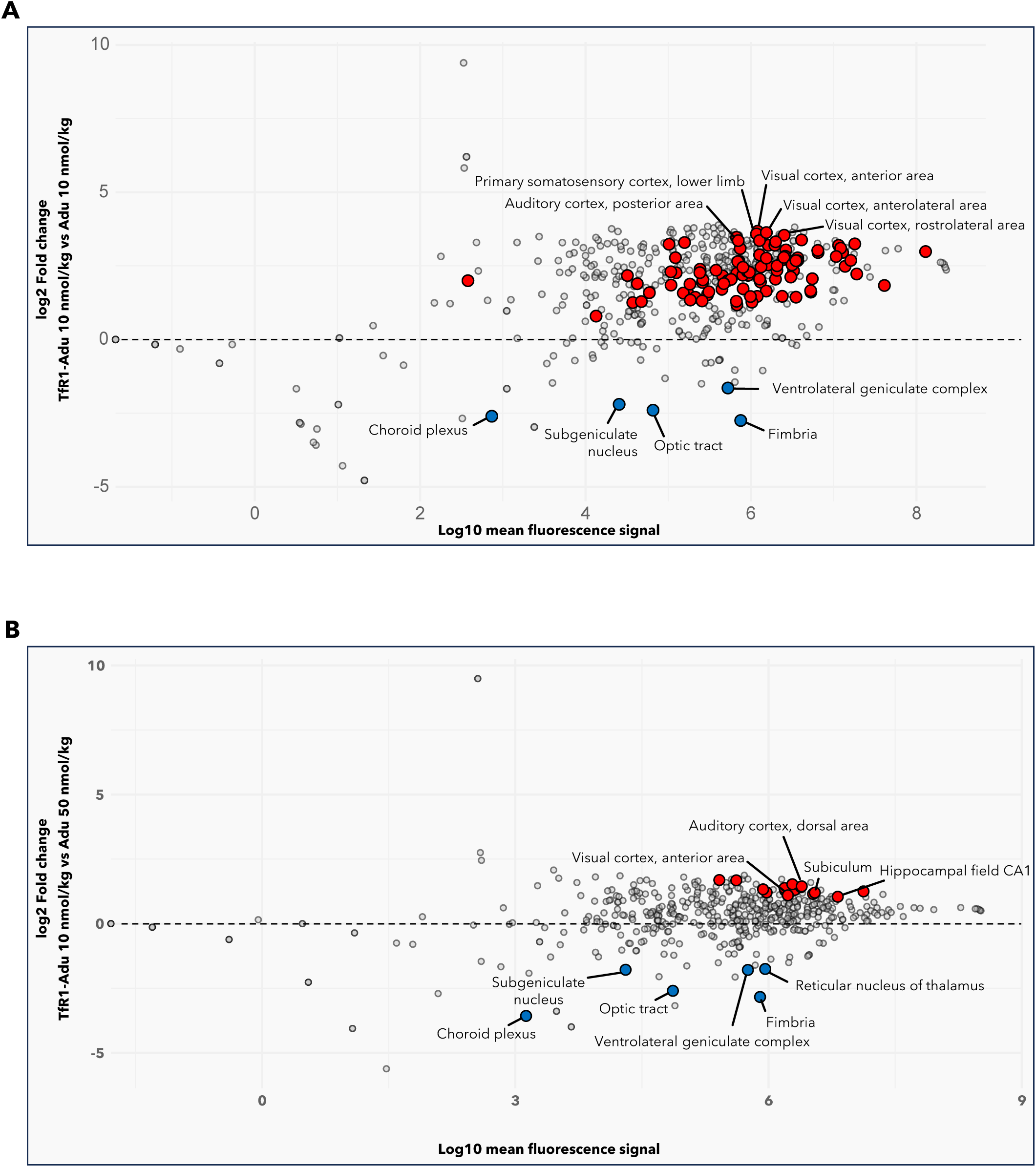
Differential brain regional levels of TfR1-Aducanamab and Aducanumab after chronic dosing in transgenic APP/PS1 mice. Volcano plots depicting brain regions with significantly higher (red dots) or lower (blue dots) levels (mean fluorescent signal) of TfR1-Aducanumab (TfR1-Adu) compared to Aducanumab (Adu). **(A)** Comparison of low-dose TfR-Adu (10 nmol/kg) vs. low-dose Adu (10 nmol/kg). **(B)** Comparison of low-dose TfR-Adu (10 nmol/kg) vs. high-dose Adu (50 nmol/kg). Grey dots represent brain regions with no significant difference in the levels of TfR1-Adu and Adu. Selected brain regions are indicated. Dunnett’s test negative binomial generalized linear model.

**Figure S4.**
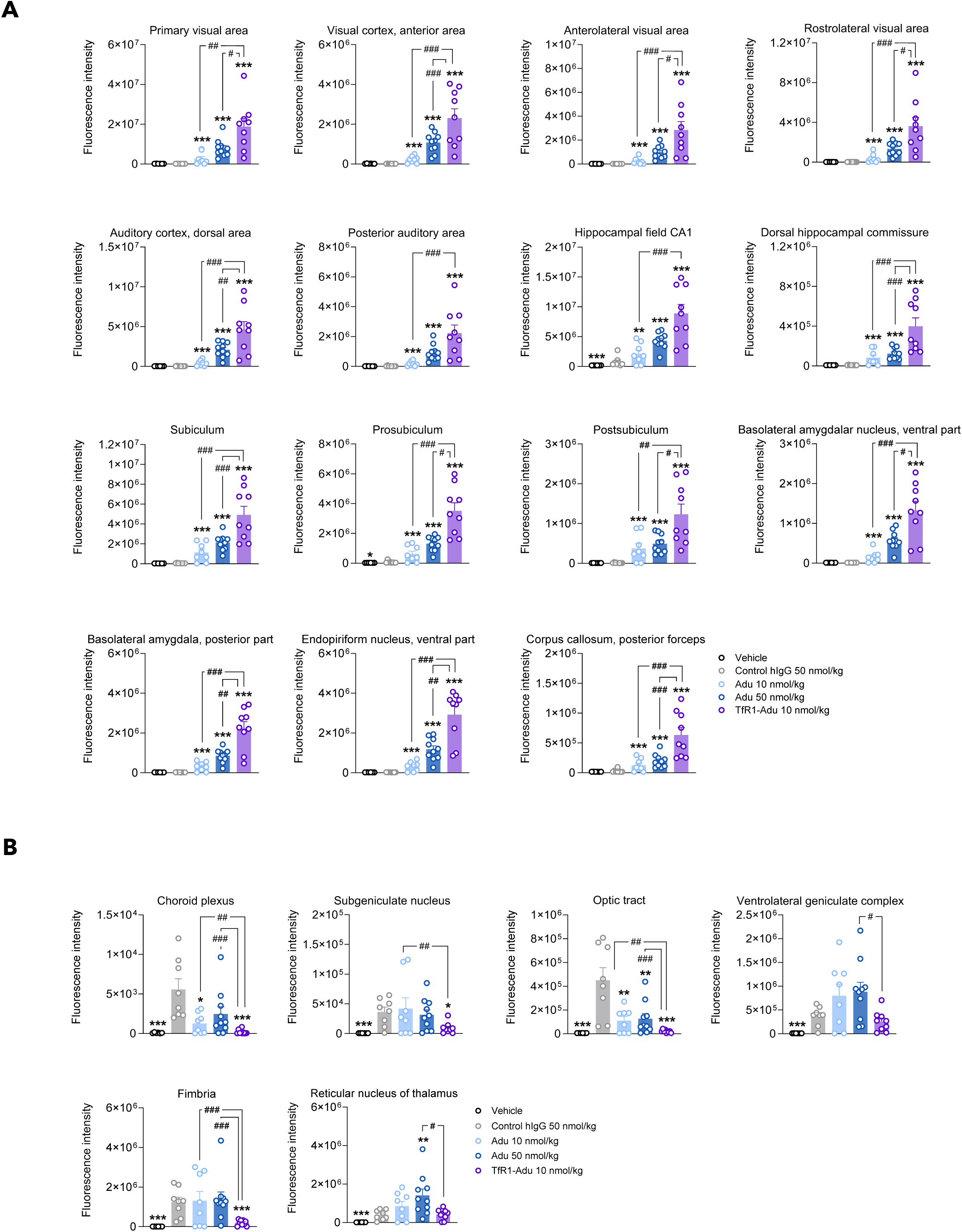
Differential brain regional levels of TfR1-Aducanamab and Aducanumab after chronic dosing in transgenic APP/PS1 mice. **(A)** Brain regions with significantly higher levels (fluorescence intensity) of low-dose TfR1-Aducanamab (TfR1-Adu, 10 nmol/kg) compared to low-dose or high-dose Aducanumab (Adu, 10 nmol/kg, 50 nmol/kg). **(B)** Brain regions with significantly lower levels (mean fluorescence signal) of TfR1-Adu (10 nmol/kg) compared to Adu (10 nmol/kg, 50 nmol/kg). *p<0.05, p<0.01, **p<0.001 vs. Control hIgG (50 nmol/kg); ^#^p<0.05, ^##^p<0.01; ^###^p<0.01 vs. TfR1-Adu. Dunnett’s test negative binomial generalized linear model.

**Figure S5.**
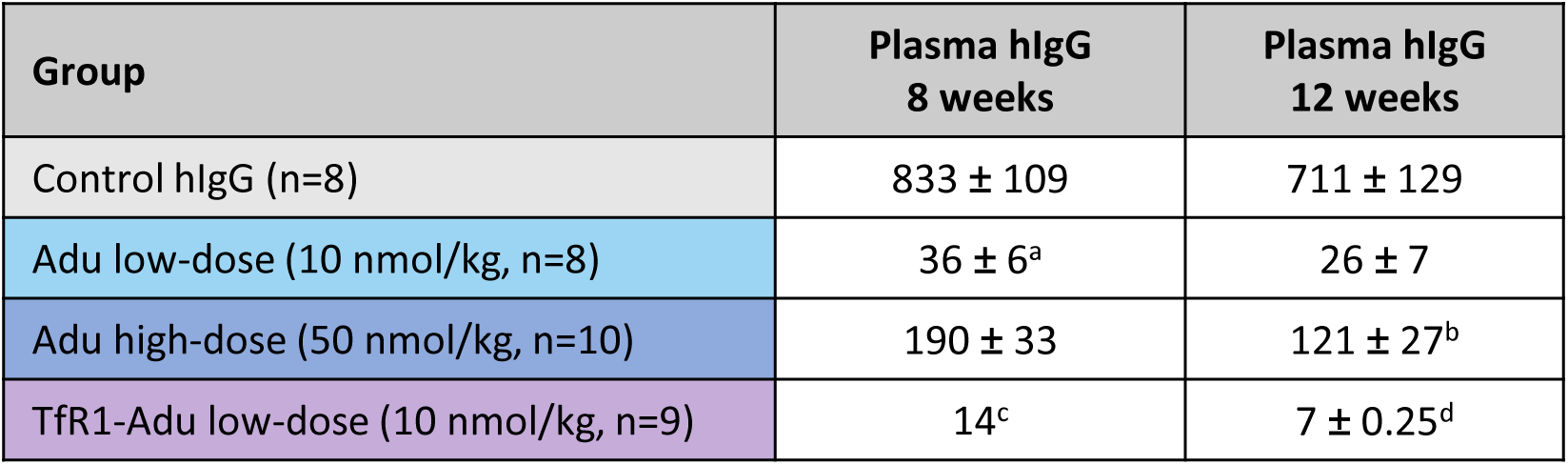
Systemic exposure of Aducanumab and TfR1-Aducanumab during and after chronic dosing in transgenic APP/PS1 mice. Aducanumab (Adu) and TfR1-Aducanumab (TfR1-Adu) exposure was determined in plasma samples in treatment week 8 (tail microbleed, 72 hrs post-dosing) and treatment week 12 (terminal heart blood, 72h after the last dose). Plasma drug concentrations are indicated as mean ± S.E.M. ^a^Adu levels were below lower level of quantification (LLOQ) in 2 out of 8 mice; ^b^Adu levels were below LLOQ in 1 out of 10 mice; ^c^TfR1-Adu plasma levels were below LLOQ in 9 out of 9 mice; ^d^TfR1-Adu plasma levels were below LLOQ in 8 out of 9 mice.

**Figure S6.**
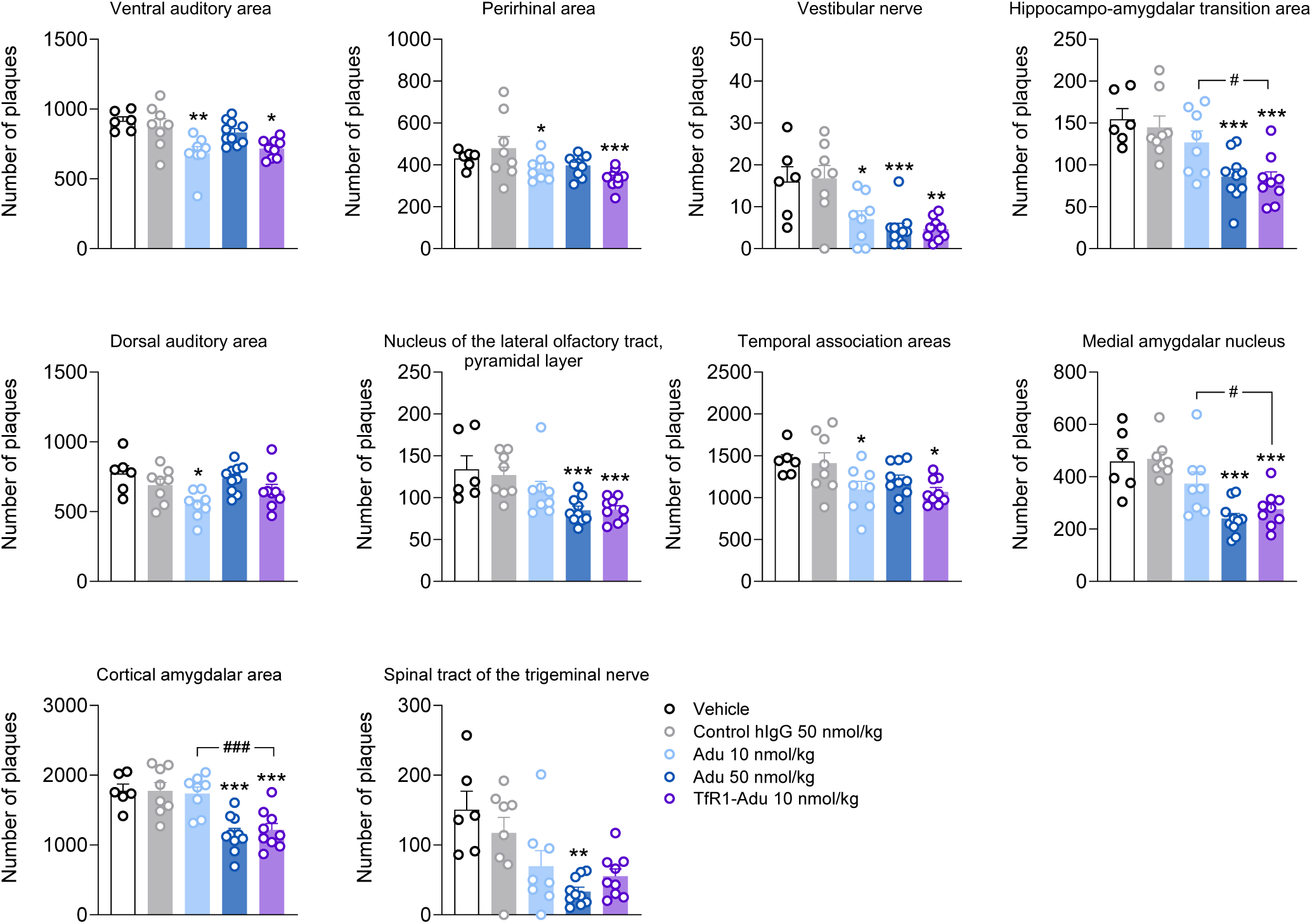
Top-10 brain regions with most pronounced reduction in β-amyloid plaque load after chronic dosing with TfR1-Aducanumab (TfR1-Adu) or Aducanumab (Adu) in transgenic APP/PS1 mice. Plaque load was determined by whole hemibrain labelling with anti-hAβ antibody. *p<0.05, **p<0.01, ***p<0.001 vs. Control hIgG; ^#^p<0.05, ^###^p<0.01 vs. low-dose TfR1-Adu (Dunnett’s test negative binomial generalized linear model).

**Figure S7.**
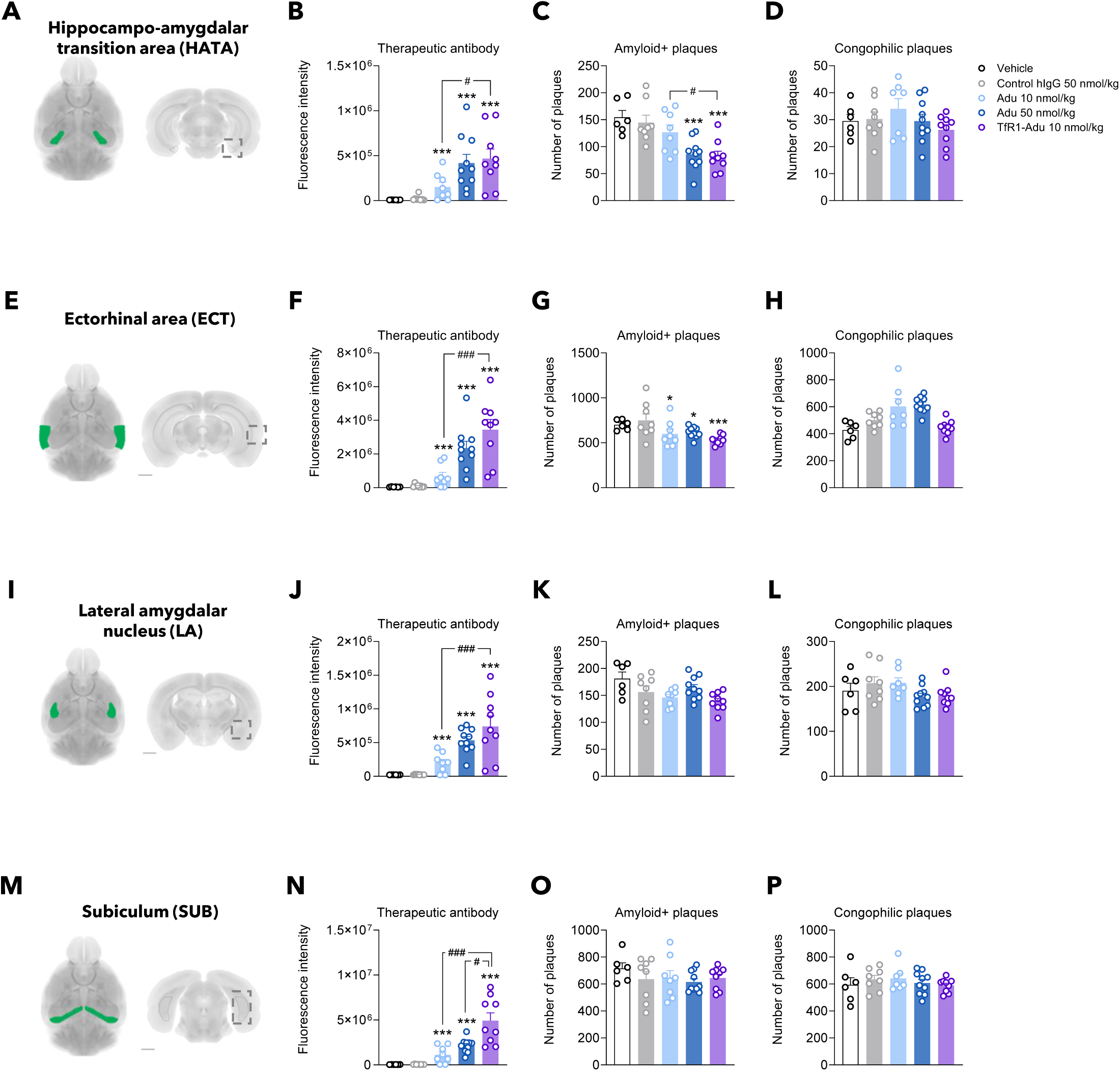
Comparison of hAβ antibody biodistribution and plaque load in selected brain regions following chronic treatment with TfR1-Aducanumab or Aducanumab in transgenic APP/PS1 mice. Chronic treatment study. **(A-D)** Hippocampo-amygdalar transition area (HATA), **(E-H)** Ectorhinal area (ECT), **(I-L)** lateral amygdala (LA) and **(M-P)** Subiculum (SUB). HATA, ECT, LA and SUB represent selected brain areas with markedly higher IgG signal of low-dose TfR1-Aducanumab (TfR1-Adu, 10 nmol/kg) compared to low-dose aducanumab (Adu, 10 nmol/kg). TfR1-Adu reduced Aβ-positive plaque counts in the HATA and ECT, but not LA and SUB. Note that TfR1-Adu or Adu did not reduce congophilic plaque counts in these four regions. The left hemibrain was co-stained with anti-human IgG (therapeutic antibody distribution, accumulated fluorescence signal intensity) and anti-human Aβ (Amyloid+ plaques). The right hemibrain was stained with Congo Red (Congophilic plaques). *p<0.05, ***p<0.001 vs. Control hIgG (50 nmol/kg); ^#^p<0.05, ^###^p<0.001 for indicated group comparisons. Dunnett’s test negative binomial generalized linear model).

**Figure S8.**
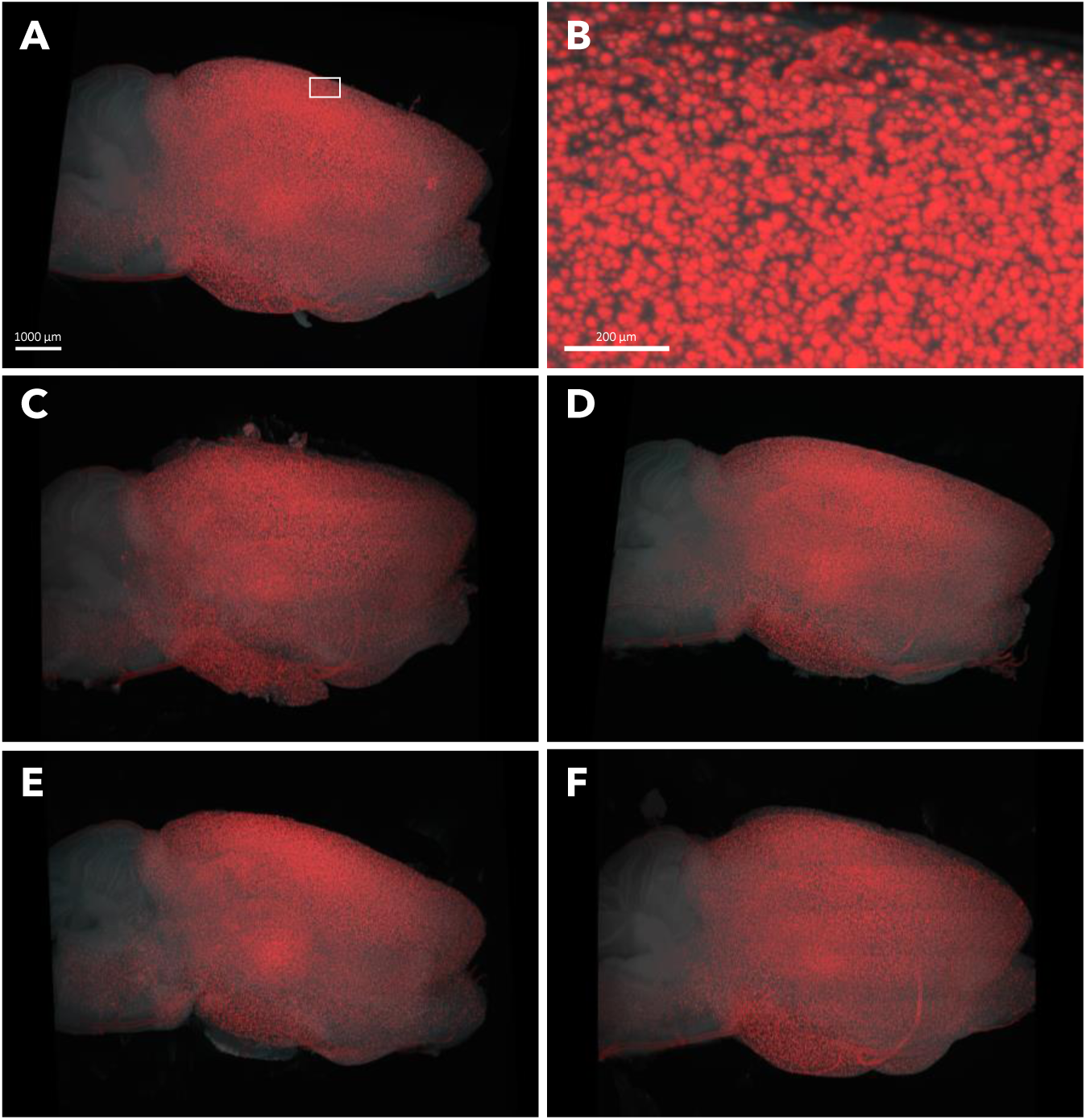
3D imaging of congophilic amyloid plaque deposition. Chronic treatment study. Representative parasagittal digital micrographs of Congo Red-stained hemibrains of APP/PS1 mice administered **(A)** vehicle (framed area further magnified in panel **B**; scale bar, 200 µm), **(C)** Ctrl-hIgG (50 nmol/kg), **(D)** low-dose Aducanumab (10 nmol/kg), **(E)** high dose Aducanumab (50 nmol/kg), or **(F)** low-dose TfR1-Aducanumab (10 nmol/kg) once weekly for 12 weeks. Scale bar, 1000 μm.

